# *PTPN11*/SHP2 negatively regulates growth in breast epithelial cells: implications on tumorigenesis

**DOI:** 10.1101/2020.07.30.228445

**Authors:** Madhumita Chakladar, Madhumathy G Nair, Jyothi S Prabhu, T S Sridhar, Devaki Kelkar, Madhura Kulkarni, LS Shashidhara

## Abstract

PTPN11/SHP2, a non-receptor protein tyrosine phosphatase is a prominent target of the receptor tyrosine kinase that participates in positive feedback signalling of the human epidermal growth factor receptors and helps in growth and migration. PTPN11/SHP2 is widely believed to be an oncoprotein, although it’s possible tumor-suppressor role is also reported. Our analysis of breast cancer metadata shows, PTPN11/SHP2 copy number loss in luminal A subtype is correlated to poor disease-specific survival and late-stage cancer at diagnosis. Analysis of the level 4 Reverse Phase Protein Array (RPPA) data available on the TCGA database resulted in positive correlations between the lower expression levels of constitutively active variant, the phospho-SHP2-Y542, of PTPN11/SHP2 and larger tumor size and lymph node positivity. We experimentally examined possible negative regulation of growth by PTPN11/SHP2 using MCF10A, a normal breast epithelial cell line. Knock-down of PTPN11/SHP2 resulted in increased cell migration, cell shape changes to mesenchymal morphology, and increased survival in cells treated with epirubicin, a DNA-damaging drug. However, it did not alter the rate of cell proliferation. It is possible that PTPN11/SHP2 might function as a tumor suppressor by potentiating proliferating cells with increased cell migration and resistance to apoptosis.

**Statement of Significance:** Molecules like *PTPN11*/SHP2, among many others that show dual specificity in tumorigenesis in the same tissue depending on the upstream signaling cues, present challenges in the field of targeted drug therapy. This study puts forth the importance of understanding the mechanism of one of the two outcomes and thereby helps better clinical management of a subgroup of cancer.

## Introduction

Src homology phosphatase 2 (SHP2) is encoded by the PTPN11 gene. PTPN11/SHP2 has two N terminal SH2 domains, N-SH2 and C-SH2, a middle phosphatase domain, and a C terminal proline-rich tail with tyrosine 542 and 580 which are phosphorylated for catalytic activation of the phosphatase by Src Kinases (Neel, Gu, and Pao 2003; Zhao, Sedwick, and Wang 2015). Characterization of phosphorylation profile regulated by PTPN11/SHP2 activity and its localization has identified 53 different proteins including ERK, P38, and many adhesion kinases (Corallino et al. 2016).

PTPN11/SHP2 is one of the few phosphatases demonstrated to have oncogenic properties. Over-expression of PTPN11/SHP2 is shown to correlate to aggressive clinical manifestations of gastric cancer (Kong et al. 2017), hepatocellular carcinoma (Han et al. 2015), laryngeal carcinoma (Gu et al. 2014), small cell lung cancer (Yang et al. 2017), thyroid cancer (Hu et al. 2015), ovarian cancer (Hu et al. 2017), glioblastoma (Sturla et al. 2011), colorectal cancer (Yu et al. 2011), pancreatic ductal adenocarcinoma (Zheng et al. 2016) and oral cancer (Xie et al. 2014). Its tumorigenic function is known to be associated with the activation of ERK and PI3-AKT pathways (Aceto et al., 2012 and Zhang et al., 2016). A contrasting tumor suppressor role is also reported for PTPN11/SHP2 in hepatocellular cancer (Bard-Chapeau et al., 2011) and oesophageal squamous cell cancer (Qi et al., 2017), which is mediated by the dephosphorylation of pSTAT3.

In the context of breast cancer, while PTPN11/SHP2 expression levels do not correlate to any intrinsic molecular subtype, its overexpression is associated with the poor overall survival in ER-positive-Ki67 low, Luminal A patients and ER-positive-Ki67 high, Luminal B/HER^−^patients (Muenst et al. 2013). Furthermore, PTPN11/SHP2 is reported to repress let-7 miRNAs in HER2 overexpressing breast epithelial cells (Aceto et al. 2012). Increased co-expression of PTPN11/SHP2 and EGFR is also correlated to basal-like and triple-negative breast cancer (Matalkah et al. 2016). Breast cancer is one of the well-studied cancers, both at the clinical and the molecular levels. Two major cancer databases, the METABRIC (specific to breast cancer) and TCGA, both house high quality datasets for breast cancer. Using those databases and experimental validation, here we report our results of the study exploring possible tumor suppressor role for PTPN11/SHP2 in the context of breast cancer.

In contrast to earlier reported oncogenic role in breast cancer, our analysis of breast cancer meta data using both TCGA and METABRIC databases indicate a tumor-suppressor role for PTPN11/SHP2. We further experimentally validated this observation using MCF10A, a non-transformed breast epithelial cell line. siRNA mediated knockdown of PTPN11/SHP2 in MCF10A promotes hallmarks of cancer like increased cell migration and decreased chemosensitivity to epirubicin, although it did not affect cell proliferation.

## Results

### Clinical Correlations of PTPN11/SHP2 copy number changes in Luminal A Subtype

We examined the correlation of PTPN11 copy number status with clinical parameters across the PAM50 subtypes of breast cancer (Bernard et al. 2009). We did not include a subset of patients with claudin-low status for our analysis as it was not defined in the PAM50 classification. We observed a significant association of PTPN11/SHP2 to certain clinical parameters within the Luminal A subtype in the METABRIC database. Copy number loss for PTPN11/SHP2 was correlated to late-stage cancer (Figure 1A) and nodal positivity (Figure 1B), but not to the age of patients at diagnosis (Fig. 1C), or their tumor size (Fig. 1D) and grade (Fig. 1E). More importantly, copy number loss was significantly associated with the poor disease-free survival (Fig. 1F) in Luminal A subtype of the breast cancer.

**Figure 1:**
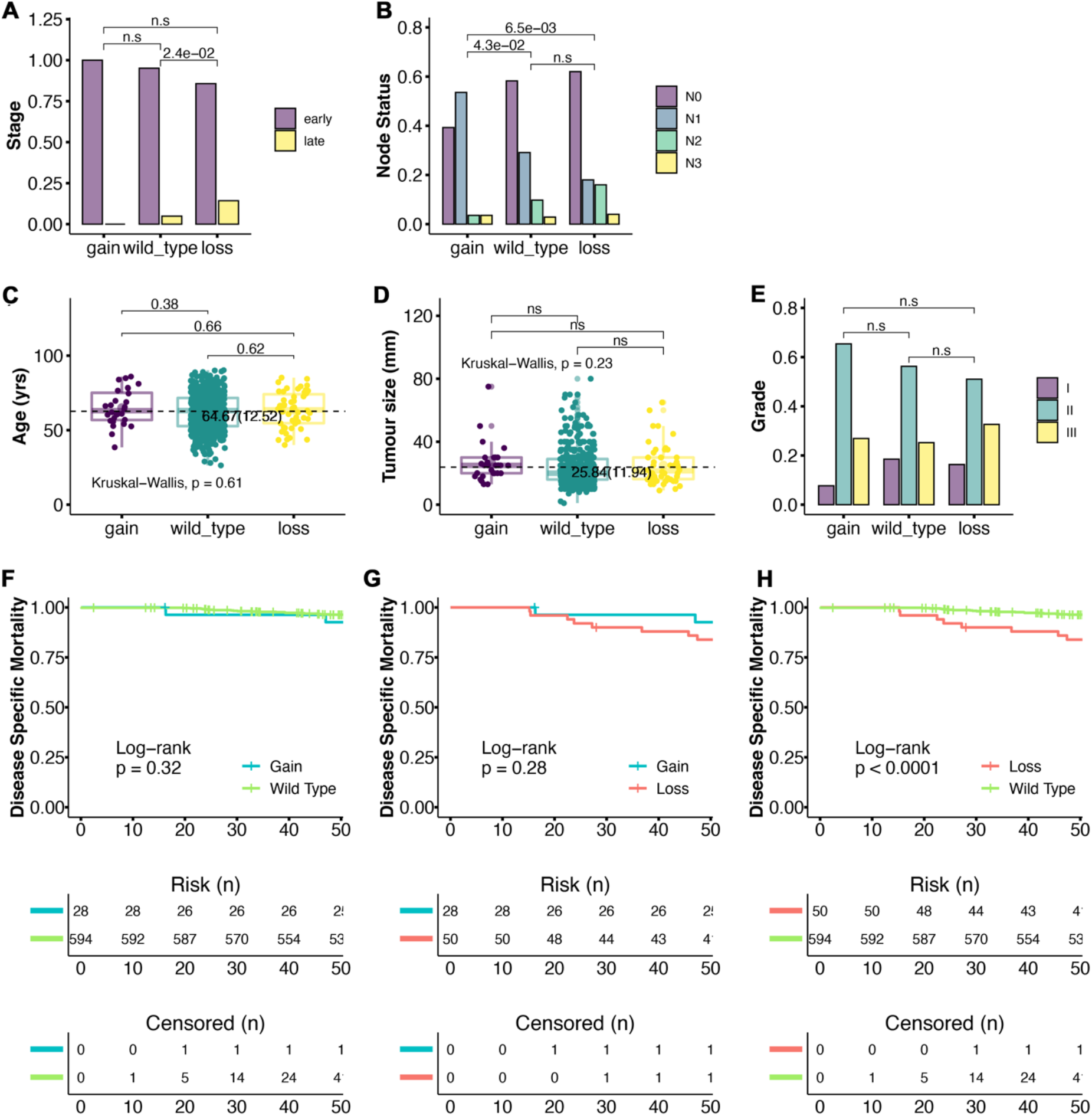
Clinical association of PTPN11/SHP2 copy number changes in Luminal A Subtype of breast cancer. Correlation between the Copy number of PTPN11 and certain clinical parameters were analyzed using the data available in the METABRIC database on Luminal A subtype of breast cancer. (A-E) Statistically significant correlation was observed for late-stage cancer (A) and nodal positivity (B), but not to the age of patients at diagnosis (C), tumor size (D) or the grade (E). In case of nodal positivity, protective role of high copy number of PTPN11 was obvious when compared to the wildtype or the loss in the copy number. (F-H) Correlations between Copy number differences between gain vs wt (F), gain vs low (G) and low vs wt (H) for the disease-free survival at 4 years of follow up. Loss of copy number is significantly associated with poor disease-free survival (H).

### Clinical Correlation of Phospho (tyrosine 542) PTPN11/SHP2 protein expression in Luminal A subtype

To re-confirm our gene-level observation described above at the protein level, we examined if the correlations between phospho-SHP2 (functionally active form of the protein) and the clinical parameters using the proteome data available in the TCGA breast cancer database. We grouped Luminal A breast cancer patients into two groups, those expressing high (above 3^rd^ quartile) and low (below 1^st^ quartile) levels of phospho-SHP2. Lower levels of expression of phospho-SHP2 protein correlated to larger tumor size (Fig. 2A) and nodal positivity (Fig. 2B), while there was no significant correlation to age (Fig. 2C) of the patients at first diagnosis or metastasis (Fig. 2D) and stage (Fig. 2E) of the cancer. Based on these two observations, we hypothesized that PTPN11/SHP2 may function as a tumor suppressor possibly under specific contexts, perhaps those that are associated specifically with the Luminal A subtype of breast cancer.

**Figure 2:**
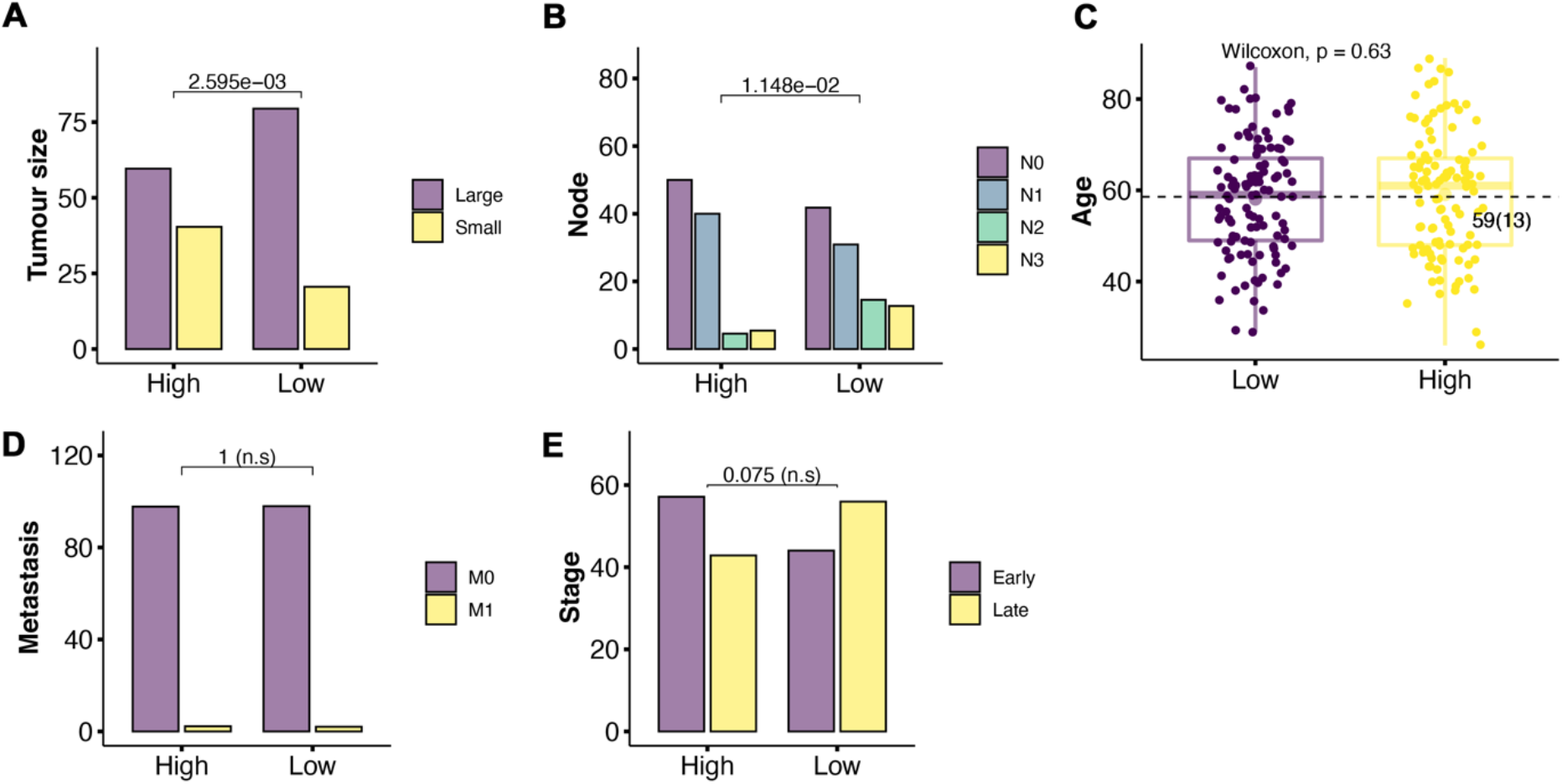
Clinical Correlation between Phospho-PTPN11/SHP2 protein expression and clinical phenotypes of Luminal A patients at diagnosis. We used proteomic data from the TCGA database to examine the correlations between the functionally active phosphor-PTPN11/SHP2 to the clinical manifestations of Luminal A patients of breast cancer. Significant association is observed between Low levels of Phospho- PTPN11/SHP2 and larger tumor size (A) and LN2 and LN3 positivity (B). No such correlations were observed for age (C), metastasis (D) or the stage of the cancer (E).

### Effect of the knockdown of PTPN11/SHP2 on the proliferation of MCF10A cell line

Knockdown of PTPN11/SHP2 in luminal A cell line, such as T47D, has been reported to decrease migration rate and EMT (Sun et al. 2017), while inhibition of PTPN11/SHP2 in luminal B cell line, MCF7, decreases cell growth (Li et al. 2014). This is in accordance with the oncogenic function of PTPN11/SHP2. However, as described above we observed that lower levels of phospho-SHP2 are associated with larger tumor size in luminal A subtype of breast cancer patients. We sought to verify possible tumor suppressor role of PTPN11/SHP2 by studying the effects of its loss in the transformation of, otherwise, normal breast epithelial cells. Using normal cells was important to examine the molecular context of its tumor suppressor role unperturbed by hormonal receptor signaling. We silenced PTPN11/SHP2 in MCF10A, a non-transformed breast epithelial cell line. We successfully knocked down PTPN11/SHP2 in the monolayer culture of MCF10A with two independent siRNAs (hereafter referred as PTPN11/SHP2#si18 and PTPN11/SHP2#si19). We achieved 50-70% depletion of SHP2 protein (Suppl. FigS1A, B) and 80-90% depletion of PTPN11 mRNA expression (Suppl. FigS1C) in transfected MCF10A cells. First, we examined effects on proliferation and survival of MCF10A cells. We used the proliferation marker Ki67 to monitor rate of cell proliferation. We did not observe any significant change in the proliferation index of MCF10A cells due to the knock-down of PTPN11/SHP2 (Fig. 3A, B). We also examined degree of survival of PTPN11/SHP2-depleted cells by MTT assay. We measured cell survival at 72 hours (Day 0), 96 hours (Day 1), 120 hours (Day 2), 144 hours (Day 3) of first transfection. We did not observe any effect on the survival of non-transformed MCF10A breast epithelial cells (Fig. 3C, D).

**Figure 3:**
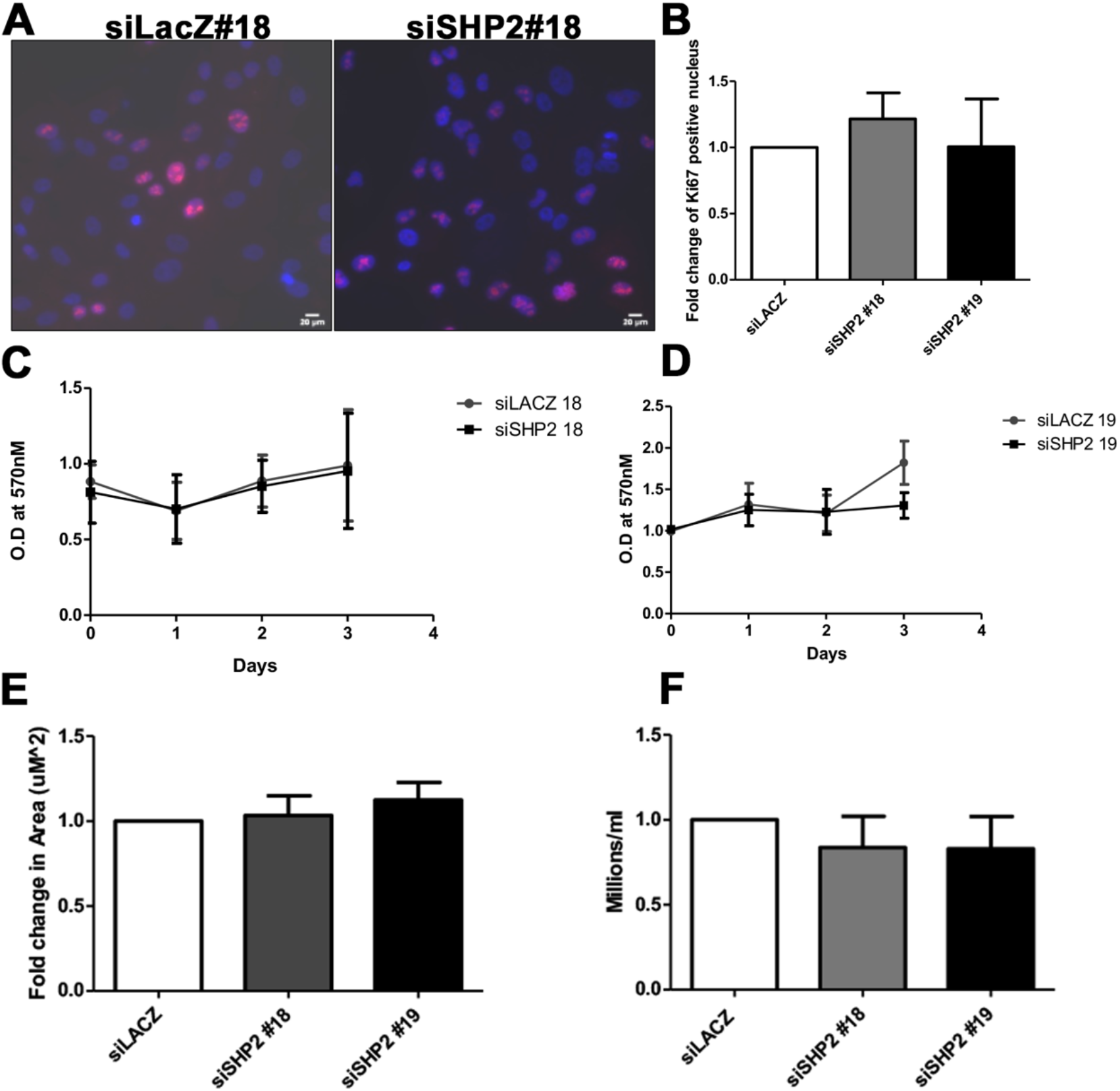
Effect of the knockdown of PTPN11/SHP2 on cell proliferation, survival, size and number. (A-B) Cell proliferation was measured in control and experimental MCF10A cells stained with the proliferation marker Ki67. Representative images of Ki67-stained cells are shown in A, both control (siLacZ#18) and the experimental (siSHP2#18). (B) shows the quantitation of change in the number of Ki67-expressing cells the control and experimental cells. We did not observe any change in Ki67 value. Images and the quantitation are shown for si18, while the observation is reconfirmed with si19 too. (C-D) Cell survival was measured using O.D values normalized to day 0 and plotted for 4 days of growth. For each biological replicate, we had 4 technical replicates. We did not observe any change in cell survival for any of the 4 days of our assays using both siRNA 18 and 19. (E) Cell size was measured by immunostaining of β-Catenin, an adheren junction protein to mark the borders of the cell. The cell size was quantitated by measuring the area of cells (area in μm^2^). At least 100 cells across 5 fields and 3 biological replicates were recorded for quantitation. There was no change observed in cell size. (F) Quantitation of cell number by Hemocytometer count. Here too we did not observe any change due to the knockdown of PTPN11/SHP2.

### Effect of PTPN11/SHP2-depletion on cell number and cell size

As lower levels of phospho-SHP2 is correlated to larger tumor size (Fig. 2A), we next examined the effect of knockdown of PTPN11/SHP2 on cell size and cell number. Here too we did not observe any effect (Fig. 3E, F). In Summary, we did not observe any significant change in any of the growth parameters of MCF10A due to the knockdown of PTPN11/SHP2 in MCF10-A cells.

### Effect of PTPN11/SHP2-depletion on cell cycle profile

PTPN11/SHP2 is known to regulate cell cycle checkpoints at G1-S transition (Tsang et al. 2012), G2-M phase (Yuan et al. 2005), and transition from metaphase to anaphase (Liu, Zheng, and Qu 2012) upon DNA damage. We did not observe any effect on the cell cycle pattern in MCF10A cells depleted for PTPN11/SHP2 (Fig. 4). We also assessed the effects of the knockdown of PTPN11/SHP2 in the background of induced DNA damage (using epirubicin: also see below). We did not observe changes in cell cycle patterns upon epirubicin treatment of MCF10A cells in the background of PTPN11/SHP2 knockdown (Suppl. Fig. S2).

**Figure 4:**
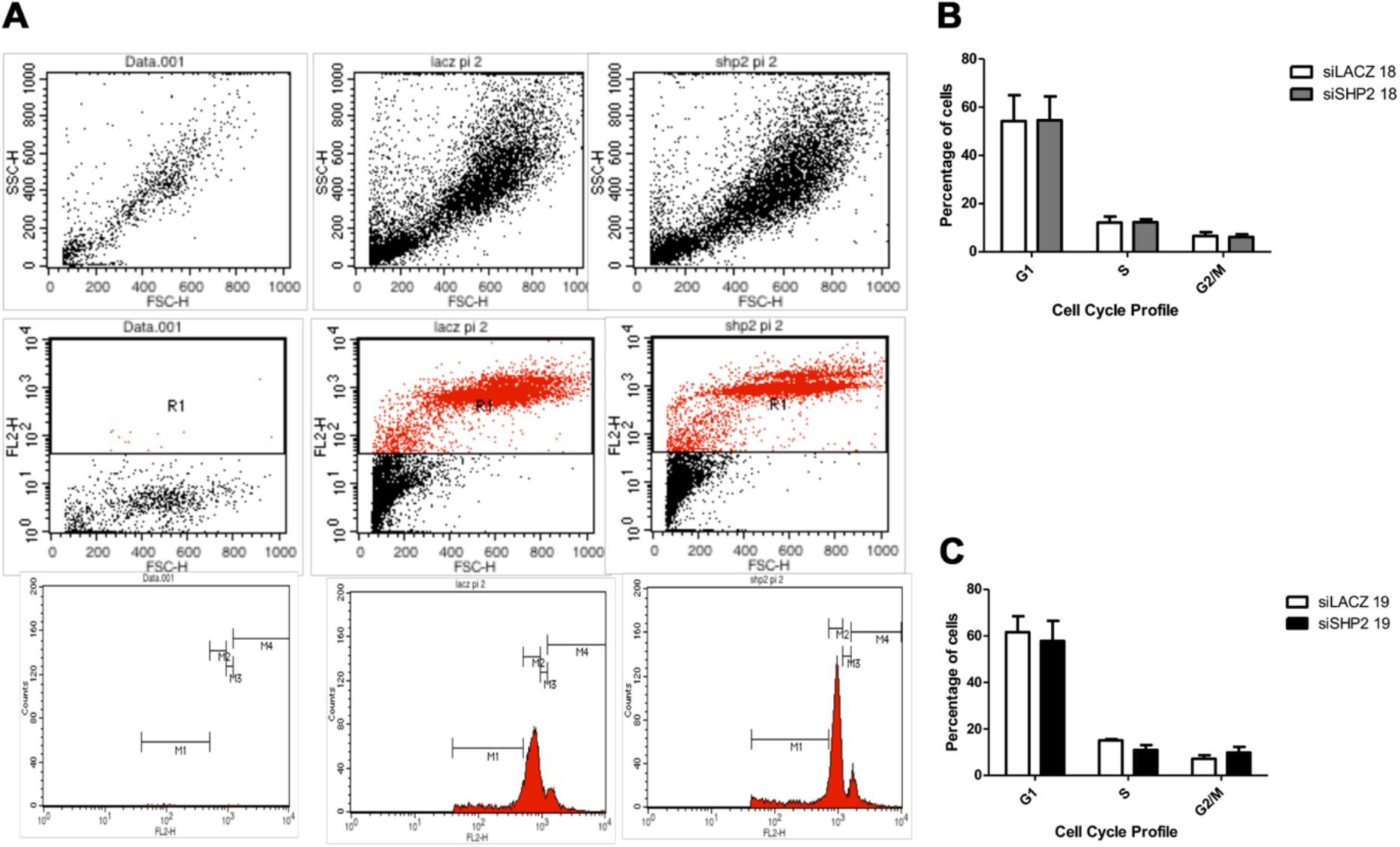
Effect of the knockdown of PTPN11/SHP2 on cell cycle profile of MCF10A cells. (A) FACS analysis showing cell cycle pattern of control and PTPN11/SHP2-depleted MCF10A cells, with scatter plots showing selected population for analysis and histogram showing cell cycle phases denoted with gate, M. Flow analysis is shown only for si18, however, data was reconfirmed with si19. M1 which is the sub G1 population was not quantitated in our analysis. For the cell cycle pattern, 10,000 cells were recorded by Flow cytometry after PI staining. (B) and (C) show quantitation of cell count in G1 (M2), S (M3), and G2/M (M4) phases of the cell cycle (N=5). No significant changes are observed for any of the cell cycle phases in PTPN11/SHP2-depleted cells.

### Effect of the knockdown of PTPN11/SHP2 on cell migration and invasion

We next assessed the effect of PTPN11/SHP2 knockdown on other hallmarks of cancer such as cell migration using the standard scratch assay and invasion using the transwell invasion assay. We observed that PTPN11/SHP2-depleted cells migrate 25% more in 24 hours of initial scratch (Fig. 5A, B). Interestingly, we also observed a change in cell morphology. Normally confluent monolayer of MCF10A cells appear cobblestone-like, whereas on PTPN11/SHP2-depleted cells acquired mesenchymal or elongated morphology (Fig. 5C). However, we did not observe any change in the invasion capacity (Fig. 5D) of PTPN11/SHP2-depleted cells in either serum-free or the media supplemented with 5% horse serum using both si18 (Figure 5E) and si19 (Fig. 5F).

**Figure 5:**
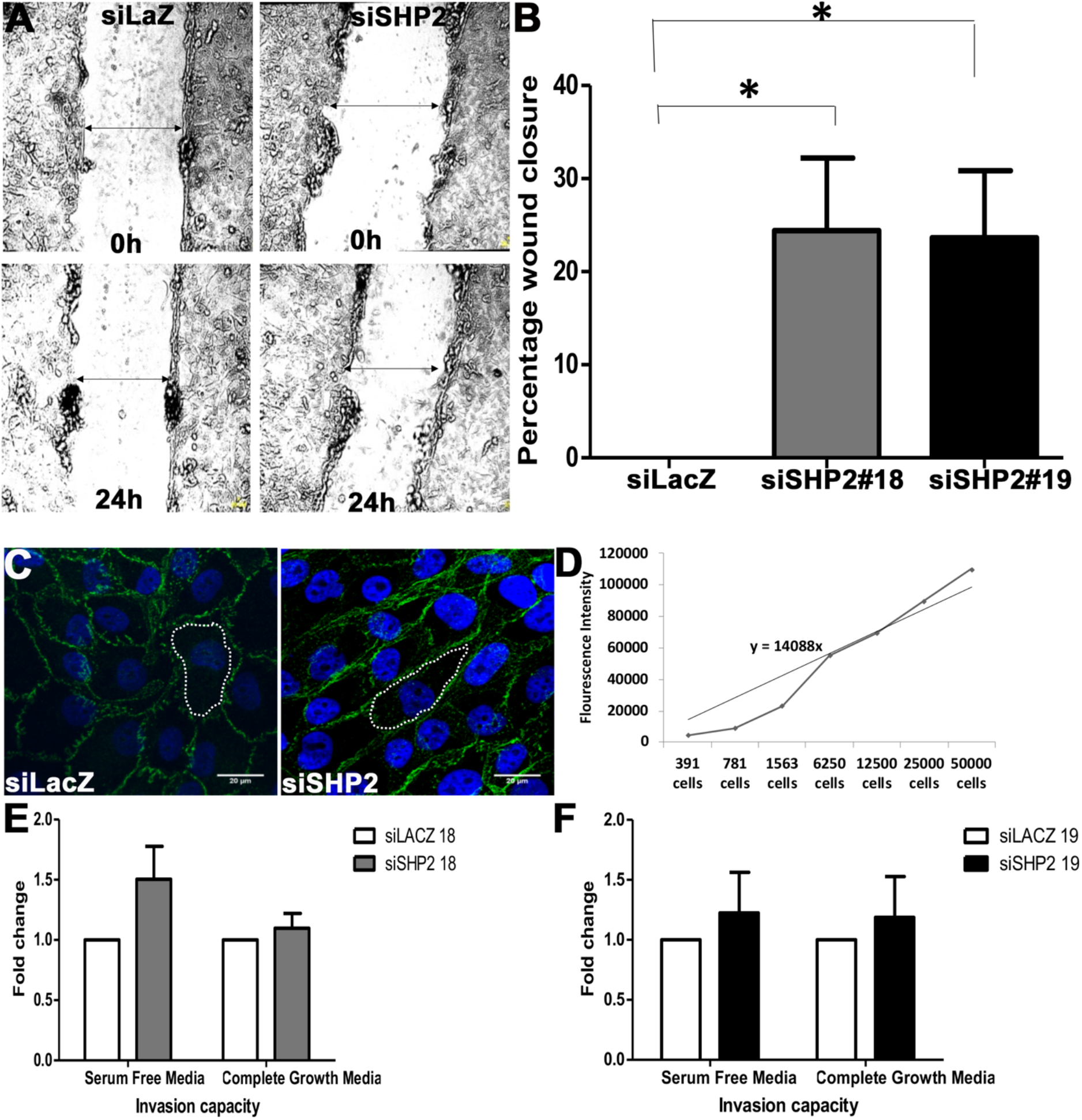
Knockdown of PTPN11/SHP2 increases the rate of cell migration and invasion, which is associated with changed cell morphology. (A) Scratch assay showing the initial wound at 0 hour and gap closure post 24 hours of scratch (shown by arrow) of normal MCF10A cells (siLacZ control) and PTPN11/SHP2-depleted cells (siSHP2). (B) Quantitation of the percentage wound closure normalized to control. Both siSHP2#18 and 19 show significantly faster would healing. (C) MCF10A cells stained with β-catenin (Green) and DAPI (Blue). White inlet used to show the change in morphology from cobblestone (in siLacZ control cells) to mesenchymal shape (in PTPN11/SHP2-depleted cells). (D) Standard curve showing fluorescence intensity versus cell number for the calibration of the transwell invasion assay Biovison kit. (E, F) The knockdown of PTPN11/SHP2 do not cause any significant changes in the invasion capacity through the matrigel coated transwell.

### Effect of the knockdown of PTPN11/SHP2 on epithelial to mesenchymal transition

We examined if the increased cell migration and the mesenchymal morphology of observed in PTPN11/SHP2-depleted MCF10A cells is an indication of epithelial to mesenchymal transition (EMT). We assessed whether depletion of PTPN11/SHP2 affects the expression of mesenchymal markers E-cadherin, N-cadherin, MMP9, Fibronectin and Vimentin. We observed approximately 2-fold increases in MMP9 (Fig. 6A), Vimentin (Fig. 6B) and Fibronectin (Fig. 6C) expression with near significance across both the siRNAs. We did not observe any change in E-cadherin levels (Fig. 6D) and could not detect N-cadherin at the desired molecular size (data not shown). We also examined the levels β-catenin by immunofluorescence, but could not detect any changes either in the nucleus or the cytoplasm (data not shown). Nonetheless, increased cell migration in the absence of any change in cell proliferation observed in PTPN11/SHP2-depleted cells could be due to increased EMT.

**Figure 6:**
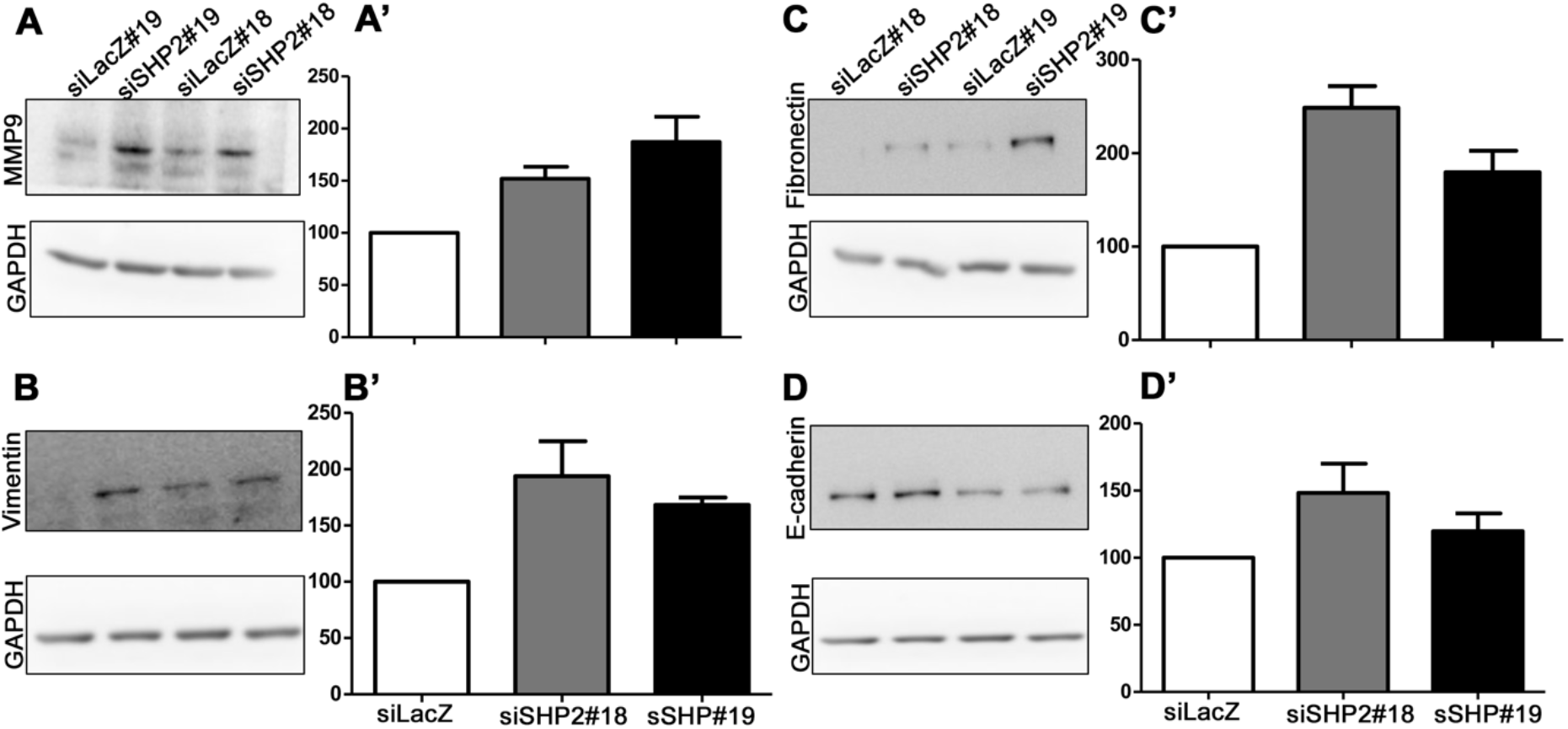
Knockdown of PTPN11/SHP2 enhances the expression of markers of epithelial to mesenchymal transition. Western blot analysis to examine the effect of the knockdown of PTPN11/SHP2 on the expression of MMP9, Vimentin, Fibronectin, and E-cadherin in MCF10A cells. GAPDH was used as loading control. Quantitation of representative blots shows a near significant increase in expression of MMP9 (A), Vimentin (B), and fibronectin (C), but no change in epithelial marker like E-cadherin (D). There were no detectable N-Cadherin levels in both control and PTPN11/SHP2-depleted cells (data not shown).

### Effect of the knockdown of PTPN11/SHP2 on the sensitivity to chemotherapeutic drugs

PTPN11/SHP2 plays a critical role in cell cycle checkpoint regulation and apoptosis upon DNA damage. PTPN11/SHP2-depletion has been reported to interfere with CHK1 activation and delay in both Cyclin E accumulation and G1-S arrest (Tsang et al. 2012). Moreover, catalytically active PTPN11/SHP2 modulates PLK1 and Aurora B activity to regulate chromosomal alignment, and thereby restoring checkpoint function at metaphase (Liu, Zheng, and Qu 2012). Alternatively, kinase-inactive PTPN11/SHP2 is involved in regulating nuclear Cdc25C translocation to the cytoplasm through 14-3-3β and inducing G2-M arrest (Yuan et al. 2005). PTPN11/SHP2 is also reported to mediate apoptosis via regulating c-ABL and caspases (Morales et al. 2014; Yuan et al. 2003). We examined the response of MCF10A cells to the treatment with DNA-damaging chemotherapeutic drug in the presence and absence of PTPN11/SHP2. We treated cells with chemotherapeutic drugs like carboplatin, epirubicin, and paclitaxel. Carboplatin inhibits replication and transcription and induces DNA breaks and cell death (Jiang et al. 2015). Epirubicin intercalates between DNA and inhibits DNA and RNA synthesis, it induces double-stranded DNA breaks and cell death (Konecny et al. 2001). Paclitaxel interferes with mitotic spindle assembly and chromosomal segregation and cell death (Jordan and Wilson 2004). We determined the Inhibitory Concentration (IC) 50 of these cell cycle and DNA synthesis interfering drugs. We used concentration in range of 100μM to 1000μM for carboplatin, 100nM-100μM for epirubicin, and 10nM-100μM for paclitaxel to optimize the IC50. We observed that MCF10A had high IC50 for carboplatin, while paclitaxel was not effective in at the concentration range tested (Suppl. Fig. S3). IC50 for epirubicin was 1μM (Suppl. Fig. S3) and this drug dose was chosen for subsequent experiments. Ability of epirubicin to introduce double-stranded DNA breaks in MCF10A cells was confirmed by measuring the levels of phosphorylated (of Serine 139) γ-H2AX (Suppl. Fig. S4).

We observed that PTPN11/SHP2-depleted cells show better survival upon epirubicin treatment at 24 hours compared to the normal cells (Fig. 7A, B). We examined if increased viability of PTPN11/SHP2-depleted cells to epirubicin treatment is associated with changes in apoptosis or cell cycle patterns. We observed that PTPN11/SHP2-depleted cells had a 2-fold decrease in early apoptotic cells (Annexin positive), late apoptotic cells (Annexin+ PI double-positive) and dead cells (PI-positive) (Fig. 7C, D). However, PTPN11/SHP2-depleted cells did not affect the ploidy or cell cycle pattern when treated with epirubicin (Supplementary Figure 2). Nonetheless, decreased chemosensitivity to epirubicin in PTPN11/SHP2-depleted cells is associated with the inhibition of apoptosis.

**Figure 7:**
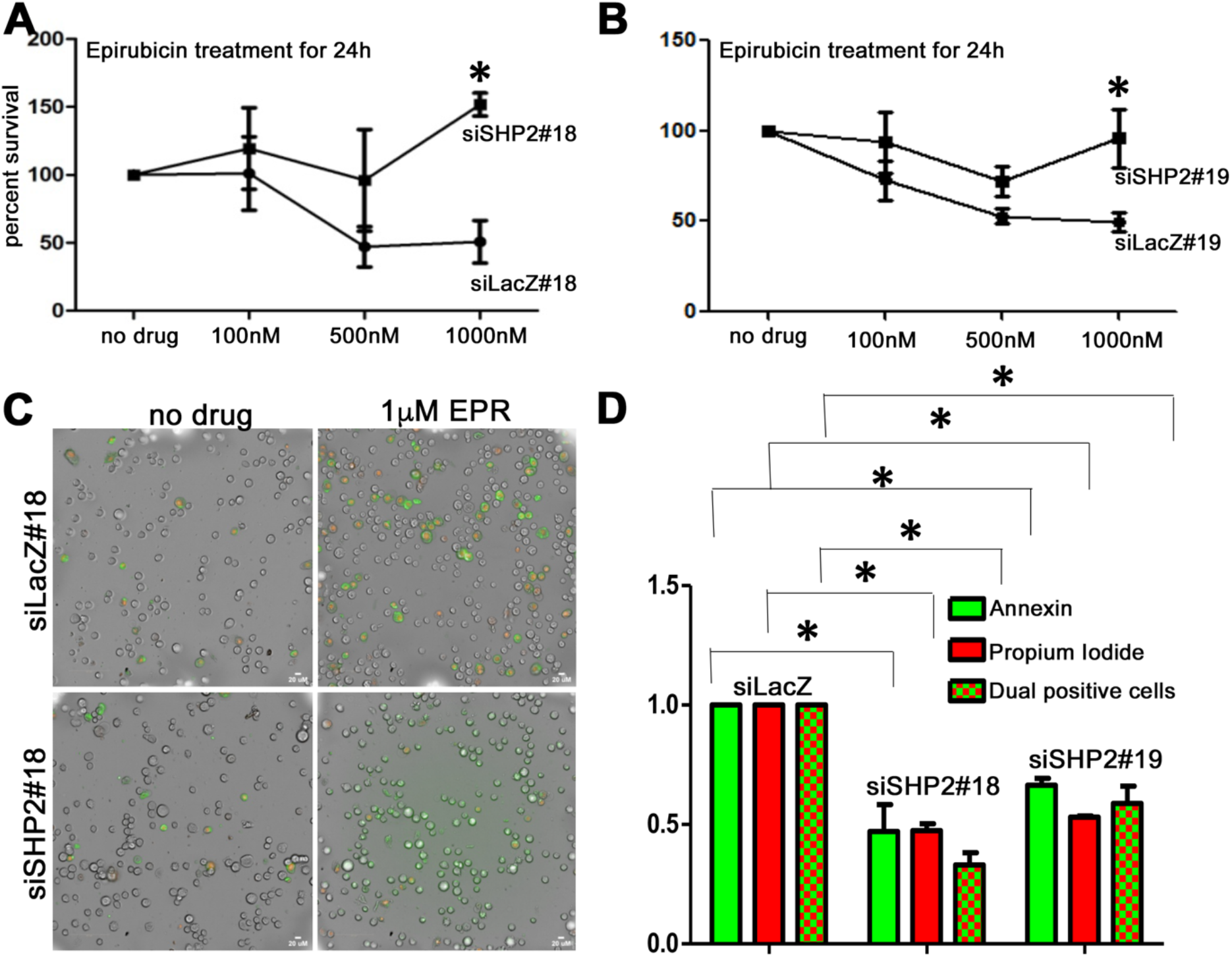
Knockdown of PTPN11/SHP2 reduces sensitivity to chemotherapeutic drugs in MCF10A cells. (A,B) MTT assay showing the dose-response curve upon epirubicin treatment in normal and PTPN11/SHP2-depleted MCF10A cells. We observed an increased survival advantage to PTPN11/SHP2-depleted cells at 1mM concentration of epirubicin. (C, D) Immunofluorescence images showing apoptosis in normal and PTPN11/SHP2-depleted MCF10A cells untreated and treated with 1mM concentration of epirubicin (C) and graphical representation of their quantitative differences (D). PTPN11/SHP2-depleted MCF10A cells show 2-fold decrease in Annexin positive, Annexin+ PI double-positive, and dead PI-positive cells.

In summary, increased cell migration, change in cell morphology, increased EMT and decreased chemosensitivity and apoptosis in response to drug treatment suggest a tumor suppressor role for PTPN11/SHP2, albeit in certain contexts.

## Discussion

PTPN11/SHP2, a non-receptor tyrosine phosphatase, participates in positive feedback regulation of EGFR pathway and drive hematologic malignancies and solid tumors including breast adenocarcinoma, prostate adenocarcinoma, lung adenocarcinoma and colorectal cancer (Aceto et al., 2012; Prahallad et al., 2015; Richine et al., 2016; Schneeberger et al., 2015; Zhang et al., 2016). In contrast to its oncogenic role, PTPN11/SHP2, through suppression of the JAK/STAT pathway, seems to function in inhibiting oncogenesis in hepatocellular carcinoma and esophageal squamous cell cancer (Bard-Chapeau et al. 2011; Qi et al. 2017). These observations suggest that PTPN11/SHP2 may function either as an oncogene or a tumor suppressor, perhaps in different cellular and molecular contexts. To verify this, we carried out a retrospective investigation of publicly available clinical metadata followed by experimental validation of our hypothesis. The oncogenic function of PTPN11/SHP2 has been reported in the backgrounds of HER2, EGFR, WNT and PI3K-AKT in breast cancer (Aceto et al. 2012; Zhao and Agazie 2015; Zhang et al. 2016).

In our analysis of breast cancer data from METABRIC and TCGA datasets, PTPN11/SHP2 does not appear to affect the clinical outcomes of the HER2-driven breast cancer or the TNBC (data not shown). However, we observed putative tumor suppressor function for PTPN11/SHP2 in Luminal A subtype of breast cancer. The copy number loss of PTPN11/SHP2 in Luminal A subtype of METABRIC cohort correlates to late-stage cancer and poor disease-specific survival. Furthermore, lower levels of expression of Phospho-PTPN11/SHP2 in luminal A correlates to larger tumor size and greater lymph node positivity in TCGA dataset. A detailed pathway enrichment analysis of phosphoproteins in TCGA RPPA Level 4 2015 data when correlated to phospho-PTPN11/SHP2 Y542 expression levels could give us insights into the context in which it may function as a tumor suppressor in the luminal A subgroup of patients in TCGA cohort. As PTPN11/SHP2 is a phosphatase, its phosphoprotein expression rather than gene expression and its implications on proteins and phospho-proteins of cancer patients is relevant in the identification of the upstream cues that is responsible for switching between its dual role in tumorigenesis. TCGA level 4 RPPA data provides the expression of a limited number of 225 protein and phosphoprotein data which could be used to understand the context of PTPN11/SHP2 function.

Experimental validation of our clinical results by transient silencing of PTPN11/SHP2 in non-transformed MCF10A showed that knockdown of PTPN11/SHP2 promotes migration, changes in cell shape to mesenchymal morphology and decreased sensitivity to chemotherapeutic drug, epirubicin, by decreasing apoptosis. Mechanistic understanding of the tumor suppressor role of PTPN11/SHP2 from our study suggests regulation of EMT molecules to limit transformation and migration of breast epithelial cells. We also report the involvement of PTPN11/SHP2 in sensitizing MCF10A cells to chemotherapeutic drugs such as epirubicin by regulating apoptosis. The phosphatase activity of nuclear PTPN11/SHP2, in response to DNA damage, in embryonic fibroblast cells activate c-ABL kinase via its SH3 domain, which in turn stabilizes P73 and allow transcription of target genes including P21^Cip1^ and BAX to initiate apoptosis (Yuan et al. 2003). PTPN11/SHP2 is also reported to mediate Rb/E2F associated apoptosis possibly by caspase8 and caspase3 activation along with PTP-1B and PTEN (Morales et al. 2014). We did not observe any change in BAX (Suppl. FigS5) or caspase 3 expression (data not shown) in PTPN11/SHP2-depleted cells with or without drug treatment. In summary, while PTPN11/SHP2 may activate apoptosis when cells are subjected to severe DNA damage (such as drug treatment), understanding the precise molecular pathway/s needs further analysis. Being a phosphatase, PTPN11/SHP2 may not behave like a classical tumor suppressor. It may function in a dose-dependent manner. Suboptimal levels above wildtype such as gain of copy number could allow it to behave differently, depending on upstream molecular cues/contexts.

## Methods

### Clinical data analysis

For all analyses, data from METBRIC 2012 and TCGA 2015 was used. METABRIC data was downloaded from cbioportal (Cerami et al. 2012) and TCGA GRCh38 data from the GDC data portal using the GenomicDataCommons R tool (Morgan M, Davis S (2019).

GenomicDataCommons:NIH/NCIGenomicDataCommonsAccess. (https://bioconductor.org/packages/GenomicDataCommons, http://github.com/Bioconductor/GenomicDataCommons.).

The TCGA RPPA level 4 data was downloaded from FireBrowse (firebrowse.org). The clinical metadata analysis was performed using R (version 3.6.1, platform x86_64-w64-mingw32/x64). Packages used for analysis include survminer_0.4.6, ggpubr_0.2.3, magrittr_1.5, survival_2.44-1.1, forcats_0.4.0, stringr_1.4.0, purrr_0.3.3 8, readr_1.3.1, tidyr_1.0.0, tibble_2.1.3, ggplot2_3.2.1, tidyverse_1.2.1, dplyr_0.8.3. For TCGA RPPA data analysis additional packages Hmisc_4.2-0, Formula_1.2-3, lattice_0.20-38 was used. Kruskal-Wallis, Wilcoxon, and log-rank test were used for statistical analysis, p<0.05 was considered significant.

### Cell culture

MCF10A cells were harvested in DMEM media (#10566-016, Thermofisher Scientific) with 100units/ml of Penstrep (#15140122). Growth media was supplemented with 5% horse serum (#26050088, GIBCO) and 20ng/ml of EGF (#E9644, Sigma), 0.5ug/ml of hydrocortisone (#H0888-5G, Sigma), 100ng/ml of cholera toxin (#C8052-1MG, Sigma) and 10ug/ml of insulin (# I1882-100MG, Sigma).

### Mycoplasma Testing

Cells were routinely checked for mycoplasma contamination and cleared (if any) using LookOutO mycoplasma elimination Kit (#MP0030).

### Cell Passaging

Monolayer MCF10A cells from passage 23 to passage 32 were used for all experiments. Media from monolayer cells was aspirated, rinsed with DPBS (3D8537-500ML), and trypsinised for 10-15 mins using 0.05% Trypsin EDTA (# 25300054, Thermofisher Scientific). The cells were incubated at 37°C, 5% CO2; dissociated cells were resuspended in DMEM with 10% horse serum and centrifuged at 2000 RPM, 6mins. Cells were seeded in a 1:4 ratio and they reach confluency of 80-90% by 3-4 days. The cells were cultured for 6 passages at any time and discarded.

### siRNA transfection

Cells were seeded at 0.16 million per 6 well and scaled down according to the plate used. 24 hours post-seeding, cells were rinsed in DPBS and grown in serum-free media (growth media without horse serum and pen strep) 24 hours before transfection. Cells were transfected using lipofectamine RNAi max (#13778150, Thermofisher Scientific) and two independent Accell siRNA-PTPN11, Targeted Region: 3’UTR (A-003947-18-0005, denoted as **#18** and A-003947-19-0010, denoted as **#19**) at 500nM and 1uM concentrations, respectively, in serum-free media. The equimolar concentration of siLACZ was used as control for each. 24 hours post-transfection, the transfection media was aspirated out and cells were replenished with growth media. Following 48 hours of transfection, growth media was aspirated out and cells were rinsed in serum-free media 1 hour before the second shot of transfection. Cells were again transfected and 48 hours post-second transfection all experiments were carried out. All experiments were carried out using both the siRNA, data for si18 is shown. Knockdown efficiency was 60-70% estimated at the protein level. The sequences for the siLACZ used are: LACZ: 5’-CGUACGCGGAAUACUUCGA-3’

3’-GCAUGCGCCUUAUGAAGCU-5’

(dTdT overhang)

### RNA isolation and RT-qPCR

Total RNA was isolated using TRIzol reagent (Sigma) and estimated using nanodrop. 500ng RNA was converted to cDNA with superscript III first-strand synthesis for RT-PCR (#1191-7010). Synthesized RNA was diluted in DNAse free water and mixed with SYBR fast qPCR master mix from Kappa biosystems (KK4601) and processed using the BioRad CFX96 real-time qPCR system. All mRNA quantification of the target gene was optimized to housekeeping control, GAPDH, or an average of housekeeping controls (ACTB, RPLPO, or PUM1) and quantitated using the ΔΔCT method or average RNU. Primer sequences used are as follows:

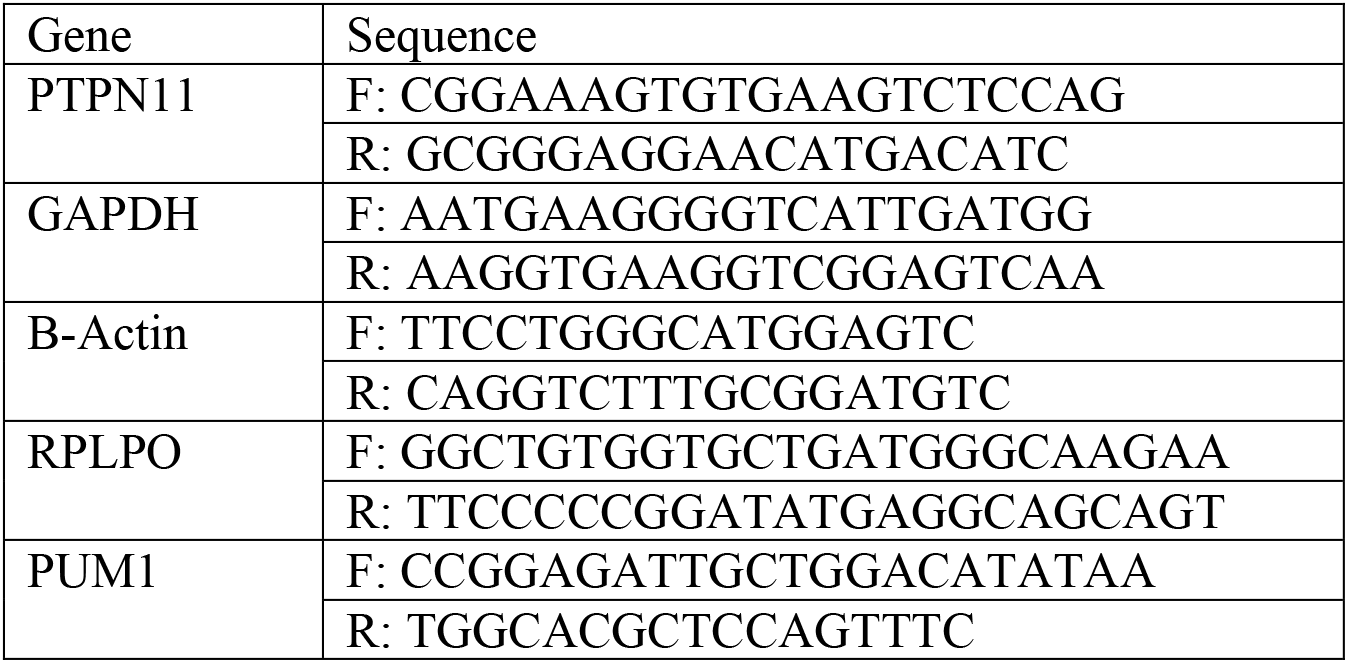

### Cell lysis and Western Blot Hybridization

Cells were washed three times in ice-cold DPBS, followed by addition of cell lysis buffer (RIPA: 20mM Tris (pH=8), 420mM NaCl, 10% glycerol, 0.5% NP40, 0.1mM EDTA, water to add up the volume) and incubated in ice for 40 mins to allow complete lysis. The lysates were collected using a cell scraper and centrifuged at 13,000 RPM, 15mins. The supernatant was collected in labelled tubes and mixed with 1X lamelli buffer and heated at 95 degrees, 5mins. SDS PAGE was run at 70V in stacking gel and at 100V in resolving gel and then transferred to the PVDF membrane for 90mins at 90V. The transferred membrane was blocked in 5% BSA or 5% Milk for 1 hour followed by the addition of primary antibody in 2% BSA or 2% Milk and then incubated overnight at 4 degrees C. Following day, the primary antibody was removed and the membrane was washed thrice with (0.1%) TBST. Secondary antibody conjugated to HRP was added in 1:10,000 dilutions in 2% BSA or 2% milk and incubated for 1hour at room temperature. Secondary antibody incubation was followed by 0.1% TBST wash and developed using an ECL kit (Merck). Densitometry analysis was used for quantitation of protein expression levels using Image J. The expression levels were normalized to housekeeping genes, GAPDH, or β Actin.

### Cell Number

PTPN11/SHP2 knocked down cells were trypsinised and centrifuged at 2000 RPM, 6mins. The cell pellets were dissolved in 1ml growth media and counted using a haemocytometer.

### Cell Size and Cell Morphology

Immunofluorescence images were captured at 63x oil objective in Leica SP8 confocal microscope. ROI of each cell was calculated for the area using Image J. A total of 100 cells across 3 biological replicates were analysed. At least 60% of the cell population were imaged and analysed for change in morphology.

### MTT assay

10ul of 5mg/ml of MTT (#M5655-100MG) was added to 100ul of cells in growth media. Growth media alone as used as blank. We incubated the cells after MTT addition for 3.5 hours at 37°C and aspirated the media with MTT. Post 3.5 hours, 100ul of DMSO (#D2438-50ml, Sigma) added, kept in a shaker for 5mins and measured absorbance at 570nM and 650nM.

### Immunofluorescence microscopy

Growth media was aspirated and cells were rinsed in DPBS. Cells were fixed with 4% PFA (Sigma) for 10mins. PFA was aspirated and cells were rinsed again in PBS, for 10mins each, repeated thrice. Cells were permeabilised and blocked with 2% FBS in 0.03% PBST (30ul Triton X (Sigma) in 10ml DPBS) for 30mins. Following permeabilization, the primary antibody was diluted in DPBS before adding and incubated overnight at 4 degrees C. Following primary incubation, cells were rinsed in 0.05% PBST (5ul Tween20 (Sigma) in 10ml DPBS) for 10mins each, repeated thrice. Cells were mounted in prolong gold Antifade DAPI or incubated in DAPI (1:1000) for 1 min and washed with DPBS before mounting (#P36931 and D9542).

### Wound healing/Scratch assay

Monolayer cells were treated with 10ug/ml of mitomycin C (Sigma M4287) for 2 hours before initial scratch. Cells were wounded using a 10ul sterile micropipette tip. Scratch was rinsed with DPBS, following which growth media were added to wells. Cells were acclimatized at 37°C for 10mins before recording a 0-hour wound distance. 3 areas per sample were recorded. 24 hours post initial wound, images of the same area recorded for 0-hour were measured with EVOS FL Auto. Wound distance was calculated using ImageJ, an average of 12 data points per sample were used for all analyses. 24-hour wound distance was subtracted from 0-hour wound distance and normalized to 0-hour wound distance (as the initial scratch was not the same across samples) and multiplied by 100 (percentage wound closure). We performed a double normalization by subtracting the percentage wound closure of every sample from its control siLACZ. Data points were plotted in GraphPad prism.

### Transwell invasion assay

K913-24 transwell assay kit was used to compare the invasion capacity of PTPN11/SHP2-depleted cells. For invasion assay, we serum-starved PTPN11/SHP2-depleted cells at 72 hours of knockdown (18-24 hours before invasion assay). Cells were then trypsinised and seeded at a concentration of 0.5-1 million cells/collagen-coated wells. 24 hours later, cells that migrated to the lower chamber were assayed using manufacturers protocol.

### Flow cytometry and cell cycle analysis

Cells were trypsinised and centrifuged at 2000 RPM, 6mins. The cell pellet was washed with DPBS by gentle vortexing and centrifuged at 2000 RPM, 6mins. The step was repeated twice. Following DPBS wash, cells were fixed in ice-cold 70% ethanol for 30 mins. Post fixation, samples were centrifuged at 2000 RPM, 6mins. The cell pellet was washed with DPBS and centrifuged again at 2000 RPM for 6mins. Cells were treated with RNAse (DS0003) (to remove any RNA contamination) for 5mins in ice. Following incubation, 5ul of Propidium Iodide in a 1million cells/sample was added 5mins before acquisition. The cell cycle profiles were acquired using BD FACS Calibur and BD FACS Celesta. Analysis was performed in BD software.

### Apoptosis

Following epirubicin treatment for 24 hours, cells in media supernatant was collected in labelled tubes. Attached cells were trypsinised and collected in the respective tube and centrifuged at 2000 RPM, 6mins, 4°C. Cells were washed with ice-cold DPBS and centrifuged at 2000 RPM, 6mins. Cell pellet was dissolved in ice-cold 90ul 1X Annexin binding buffer and added 5ul of Annexin V and 5ul Propidium iodide. The samples were incubated for 5mins with 0.25mM CaCl2 in dark and imaged and analysed in Operetta, Perkin Elmer.

### Antibodies

The following antibodies were used,

| Antibody Name | Catalog number | Working Conc. | Dilution in |
| --- | --- | --- | --- |
| SHP-2 (D50F2) Rabbit mAb | 3397 | 1:1000 | 2% BSA |
| Phospho-SHP-2 (Tyr542) Antibody rabbit mAb | 3351 | 1:1000 | 2% BSA |
| Anti-γH2A.X (phospho S139) antibody [9F3] | ab26350 | 1:10,000 | 2% Milk |
| Anti-Ki67 antibody [EPR3610] | ab92742 | 1:100 | DPBS |
| Anti-MMP9 antibody [EP1254] | ab76003 | 1:10,000 | 2% BSA |
| Purified Mouse Anti-E-Cadherin Clone 36/E-Cadherin (RUO) | 610181 | 1:1000 | 2% BSA |
| Purified Mouse Anti-N-Cadherin Clone 32/N-Cadherin (RUO) | 610920 | 1:1000 | 2% BSA |
| Purified Mouse Anti-Fibronectin Clone 10/Fibronectin (RUO) | 610077 | 1:1000 | 2% BSA |
| β-catenin | #9562S | 1:1000/1:500 | 2% BSA or DPBS |
| Propidium Iodide - 1.0 mg/mL Solution in Water | P3566 | 1ul/million cells | DPBS |
| Annexin V, Alexa Fluor® 488 Conjugate | A13201 | 5ul/million cells | DPBS |
| Vimentin | V9 clone, Santacruz | 1:1000 | 2% BSA |
| β-actin | Santacruz | 1:1000 | 2% BSA |
| GAPDH | RG000110 | 1:2500 | 2% Milk |
| Secondary Anti-Rabbit Alexa 568 | Invitrogen, Molecular probes | 1:10,000 | DPBS |
| Secondary Anti-Rabbit Alexa 488 | Invitrogen, Molecular probes | 1:10,000 | DPBS |
| Secondary Anti-Rabbit HRP | Invitrogen, Molecular probes | 1:10,000 | 2% Milk/BSA |
| Secondary Anti-Mouse HRP | Invitrogen, Molecular probes | 1:10,000 | 2% Milk/BSA |

### Statistical Analysis

Column statistics (GraphPad Prism) was used for statistical analysis, p<0.05 was considered significant. Unpaired T-Test was used for survival assay. P values are flagged as * (p<0.05), ** (p<0.01) and *** (p<0.001).

## Acknowledgements

This work was primarily supported by an Indo-Danish research grant from Department of Biotechnology, Govt. of India to LSS and TSS. JC Bose Fellowship and grant from Department of Science & Technology, Govt of India to LSS. MK is supported by Ramalingswami fellowship from Department of Biotechnology, India. We thank other members of both the laboratories for critical input.

## Conflict of interest

We declare “no-conflict-of-interest”.

## Notes

### Competing Interest Statement

The authors have declared no competing interest.

