## Supplemental Figures for "*PTPN11*/SHP2 negatively regulates growth in breast epithelial cells: implications on tumorigenesis"

**Supplementary Figures and Figure Legends**

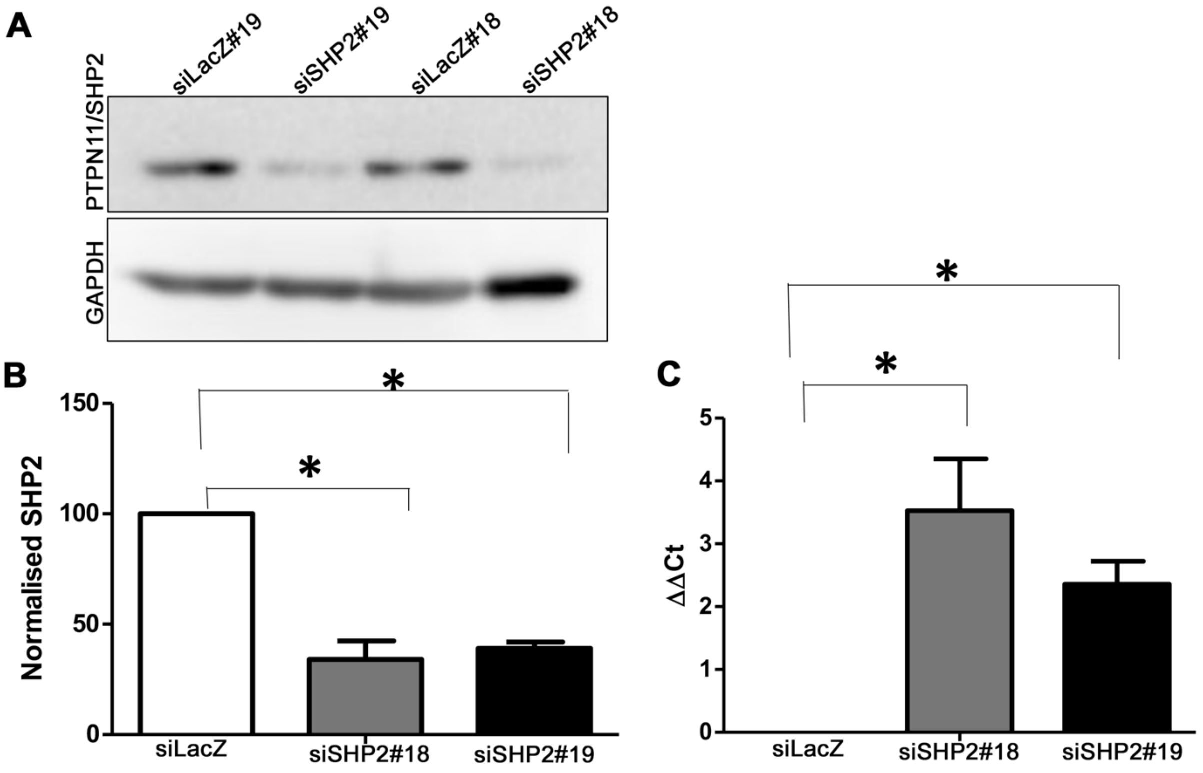


**Figure 1 siRNA mediated silencing of PTPN11/SHP2 in MCF10A cells.** Western blot (A,B) and RT-qPCR (C) anayses to assess the levels of expression of PTPN11/SHP2 protein in MCF10A cells transfected with individual siRNA (#18 and 19) against PTPN11/SHP2. GAPDH was used as loading control for Western Blot analysis. (B) Densitometric analysis of the protein expression levels show, at least, 60% knockdown. (C) RT-qPCR shows 2.5-3.5 CT difference from siLACZ control suggesting severe downregulation of PTPN11/SHP2 mRNA. RT-qPCR data was normalized to the average CT values for housekeeping gene controls, β-Actin, Pum1 and RPLPO.

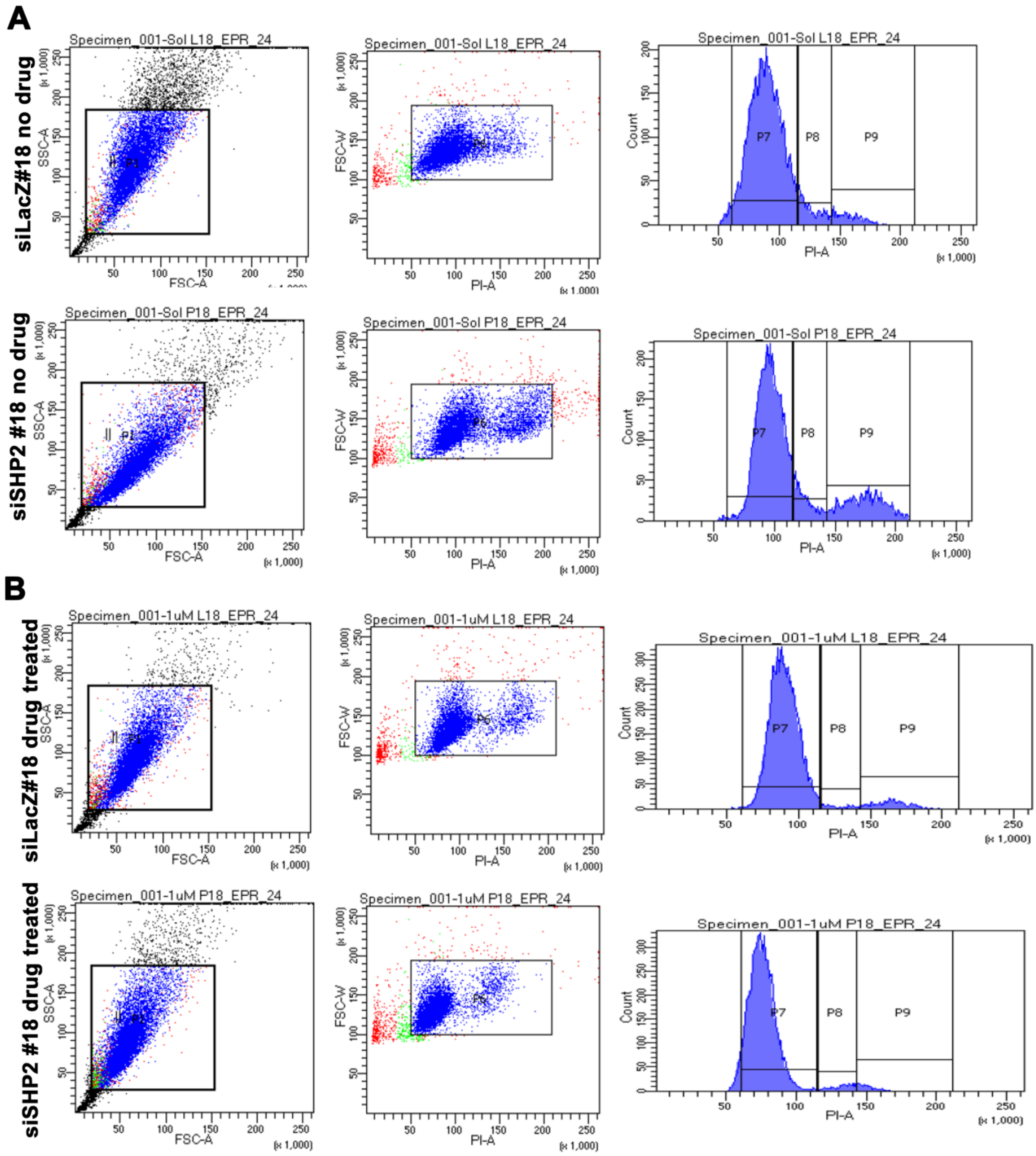


**Figure 2: FACS analyses to assess the effect of epirubicin treatment for 24 hours on the cell cycle patterns**. (A) Scatter plots showing selected population for analysis and the corresponding histograms showing cell cycle phases denoted with gate, P. (A) Flow analysis for no-drug control for both siLacZ#18 and siPTPN11/SHP2#18 MCF10A cells. (B) Flow analysis for siLacZ#18 and siPTPN11/SHP2#18 MCF10A cells treated with 1µM epirubicin (EPR). siPTPN11/SHP2#18 is shown, data was similar for siPTPN11/SHP2#19.

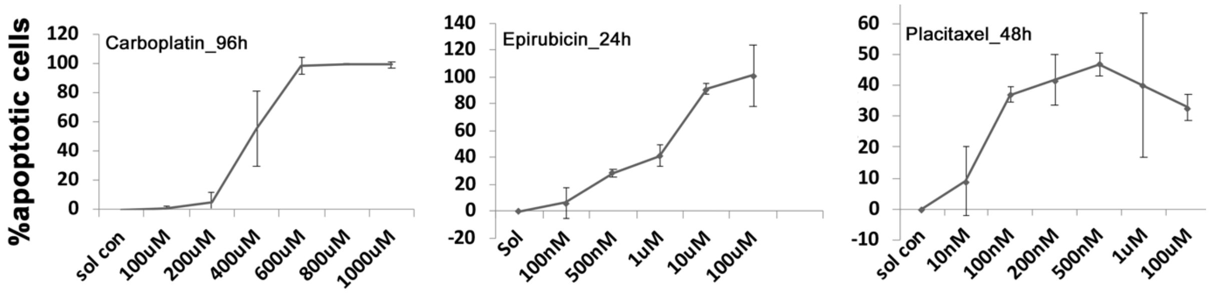


**Figure 3: MTT assay to determine dose-response curve for 3 chemotherapeutic drugs in MCF10A cells.** IC50 at 24h of treatment was 400μM for carboplatin (A) and 1μM for epirubicin (B). IC50 could not be determined for Paclitaxel as the cell death plateaued at 45-50 percent.

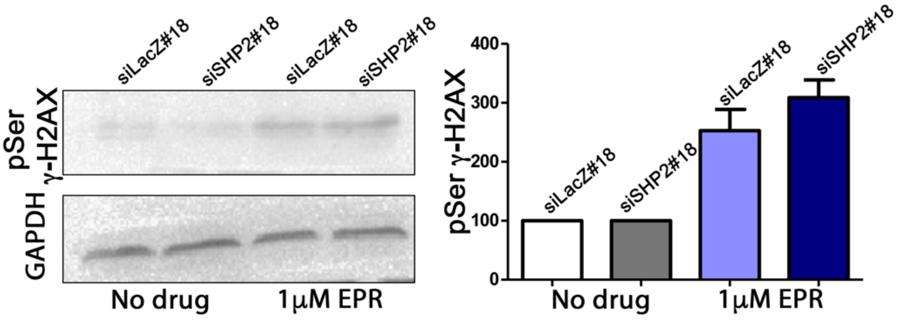


**Figure 4:** Effectivity of epirubicin (EPR), an anthracycline which induces double-stranded DNA breaks in MCF10A cells. Western blot analysis for the DNA-damage marker Phoshorylated (at Serine 139) form of γH2AX (pSerine-γH2AX). We observed significant upregulation of pSerine-γH2AX treated with epirubicin for 24hours both in normal and siPTPN11/SHP2#18 MCF10A cells. Quantitation by densitometry analysis suggests a 3-fold increase in pSerine-γH2AX.

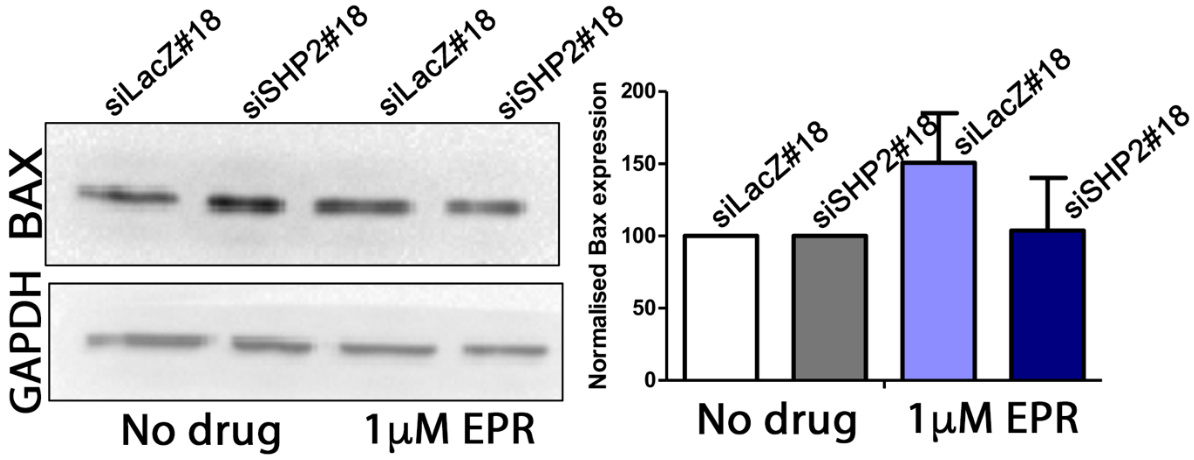


**Figure 5:** Decreased apoptosis on epirubicin-treated MCF10A cells depleted for PTPN11/SHP2 is not associated with change in the levels of BAX expression. Western blot analysis for BAX and quantitation by densitometry analysis suggests no change in BAX levels.
